# Lateral cell polarization drives organization of epithelia in sea anemone embryos and embryonic cell aggregates

**DOI:** 10.1101/2024.04.07.588493

**Authors:** Tavus Atajanova, Emily Minju Kang, Anna Postnikova, Alivia Lee Price, Sophia Doerr, Michael Du, Alicia Ugenti, Katerina Ragkousi

## Abstract

One of the first organizing processes during animal development is the assembly of embryonic cells into epithelia. In certain animals, including Hydra and sea anemones, epithelia also emerge when cells from dissociated tissues are aggregated back together. Although cell adhesion is required to keep cells together, it is not clear whether cell polarization plays a role as epithelia emerge from disordered aggregates. Here, we demonstrate that lateral cell polarization is essential for epithelial organization in both embryos and aggregates of the sea anemone *Nematostella vectensis*. Specifically, knock down of the lateral polarity protein Lgl disrupts epithelia in developing embryos and impairs the capacity of dissociated cells to epithelialize from aggregates. Cells in *lgl* mutant epithelia lose their columnar shape and have mispositioned mitotic spindles and ciliary basal bodies. Together, our data suggest that in *Nematostella*, Lgl is required to establish lateral cell polarity and position cytoskeletal organelles in cells of embryos and aggregates during *de novo* epithelial organization.

## Introduction

Epithelia emerge as polarized layers of interconnected cells in a highly divergent and context dependent manner across all animals ^1,2^. With the organization of their cells into epithelia, tissues acquire barricaded compartments and territories with often distinct developmental trajectories. During embryonic development, epithelia emerge as cells connect via lateral adhesions and differentially pattern their two opposing domains; the apical domain develops at the interface with the extracellular environment or lumen and the basal domain develops where the monolayer faces adjacent tissues or the basal membrane ^2,3^. Remarkably, in some species epithelia emerge from reaggregated cells of dissociated tissues as a first step toward rebuilding the animal body. Such whole-body reconstitution is observed in sea urchins, sea stars and cnidarians, and begins with the sorting and organization of reaggregated cells into inner and outer epithelia ^4–9^. Studies in various experimental systems pointed to differential adhesion and tension as powerful drivers of cell sorting into distinct compartments ^10,11^. However, it is not clear how dissociated cells organize into epithelia in the first place. Although intercellular adhesion is essential, whether cell polarity plays a role in this process remains unknown. It is also not clear whether embryonic and aggregate epithelialization rely on similar molecular mechanisms.

The sea anemone *Nematostella vectensis* is an excellent system to investigate these questions, as *Nematostella* epithelia emerge from both embryos and aggregates of dissociated tissues. Moreover, due to its positioning in the sister group to bilaterians, *Nematostella* is studied to provide insight into the ancestral mechanisms of animal development ^12,13^. Embryos assemble into polarized monolayers by the 16-cell stage ^14,15^. Similar to other epithelia, polarization of embryonic cells becomes evident with the differential localization of proteins in the apical and lateral domains. Conserved cadherin-catenin complexes in the apical junctions keep cells together and polarity proteins localize to their apical (Par6 and aPKC), junctional (Par3) and lateral (Par1 and Lgl) domains ^15–18^ . To investigate whether polarity is essential for *Nematostella* epithelialization, we knocked down specific polarity proteins and examined whether aggregates from dissociated mutant embryos epithelialized successfully. We found that organization of lateral polarity is crucial for epithelialization in both embryos and aggregates. Specifically, knocking down the lateral polarity protein Lgl impaired embryonic epithelia by disrupting cell shape and orientation of cell cleavages. Strikingly, *lgl* mutant epithelia exhibited a dramatic mispositioning of their ciliary basal bodies. Lgl was originally discovered in *Drosophila*, and was shown to function as a tumor suppressor by regulating both epithelial organization and cell proliferation ^19,20^. In the absence of an obvious effect on cell proliferation, our results indicate that, in *Nematostella,* Lgl helps organize embryonic and aggregate epithelia as a cytoskeletal regulator.

## Results

### Lateral polarity proteins are essential for epithelialization of *Nematostella* embryonic cell aggregates

Self-organization of tissues from aggregated cells begins with the emergence of epithelial layers. *Nematostella* aggregates form a smooth layer around their colony within 6-12 hours post dissociation (hpd). By 24hpd outer (ectodermal) and inner (endodermal) layers are defined in structure and fate identity ^8^.

We sought to examine whether distinct epithelial polarity proteins are important for *de novo* epithelialization of aggregates. To this end, we first used short RNA hairpins to knock down conserved apical and lateral polarity proteins in *Nematostella* embryos and then dissociated mutant embryos and reaggregated their dissociated cells into a solid cell mass. We hypothesized that certain epithelial proteins may be required for epithelialization after cell aggregation. In the event where dissociated cells lack a critical epithelialization protein, aggregates will fail to organize and eventually fall apart. We individually knocked down genes encoding well conserved structural polarity proteins, the apical Par3 and Par6 and the lateral Lgl and Scribbled. Previous work has shown that in *Nematostella*, Par6 localizes throughout the apical domain and is enriched together with Par3 at the apical cell junctions, while Lgl is enriched along the lateral cell contacts ^15,16^. After dissociation of treated post-gastrula embryos, we aggregated single cells via mild centrifugation as previously described ^8^. We then examined aggregate behavior in two ways: 1) By fixing aggregates at 0hpd and at 24hpd, staining for F-actin and DNA and visualizing by confocal microscopy (Figure 1). 2) And by recording aggregate behavior by live time-lapse imaging (Figure S1).

**Fig 1.**
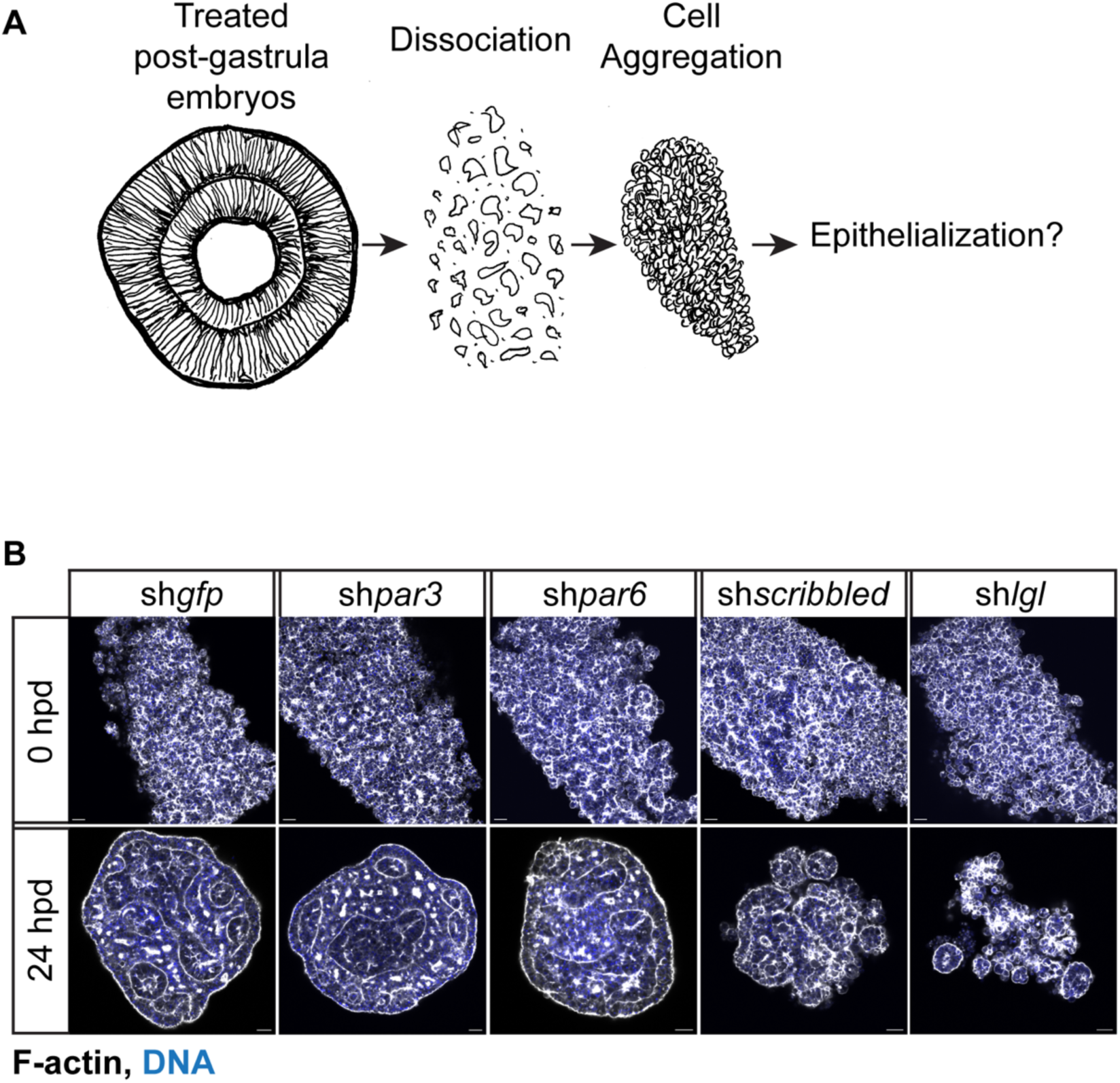
Lateral polarity proteins are essential for epithelialization of *Nematostella* embryonic aggregates. (A) Schematic of the experimental process starting with embryo dissociation and aggregate formation as described previously ^8^. (B) Aggregate formation from embryos with knocked-down expression of *gfp* (control), *par3*, *par6*, *scribbled* and *lgl*, at time of aggregation (0hpd) and 24hpd. Epithelialization manifests by organization of cells in a common aggregate with a polarized layer around multiple epithelial clusters with enclosed lumens. Cells from dissociated sh*lgl* embryos fail to organize in a common epithelial aggregate and small epithelializing clusters eventually disintegrate. Similarly, aggregates from sh*scribbled* embryos show no obvious enveloping epithelial layer, although more cells remain in contact within a common aggregate. One of four aggregates is shown from one of three independent experiments. Images show mid-sagittal sections. Scale bars: 20μm.

Aggregates from sh*par3* and sh*par6* knockdown embryos appeared to epithelialize similarly to those from control (sh*gfp*) embryos. A continuous epithelial monolayer organized around multiple internal lumens, indicating formation of both ectodermal and endodermal territories. In contrast, aggregates from sh*lgl* and sh*scribble*d embryos lacked an enveloping layer on their periphery. Notably, fewer sh*lgl* cells maintained adhesions and their aggregates thinned into small epithelial clusters that disintegrated within 48hpd (Figure 1B, Figure S1).

Live observation of aggregates by time lapse imaging confirmed that aggregates from sh*gfp* control, sh*par3* and sh*par6* embryos all compacted into rigid cell assemblies. Similar to our fixed aggregate imaging, sh*lgl* and sh*scribbled* embryos failed to compact and cell clusters from sh*lgl* aggregates disintegrated further. Interestingly, epithelialization differences across aggregates appeared as early as 5.5hpd; a critical time point by which ectodermal epithelia become apparent ^8^. By 11hpd, sh*lgl* aggregates disintegrated into progressively smaller fragments that remained separate by the end of imaging (22hpd).

### Expression of *lgl* and *scribbled* is essential for epithelial organization of developing *Nematostella* embryos

We reasoned that the lack of epithelialization in sh*lgl* and sh*scribbled* aggregates may be traced back to the compromised epithelial integrity of treated embryos. In the absence of *lgl* expression, embryos appeared to develop and gastrulate normally.

However, on closer observation of sh*lgl* embryos stained with FITC-phalloidin to outline cell boundaries, we noticed an unusual number of depressions in the ectodermal epithelia (Figure 2). While mitotic round cells are normally encountered in the apical domain of the epithelium, in sh*lgl* embryos many non-mitotic round cells were observed in the basal domain. At some regions, ectodermal depressions appeared in the vicinity of basally located round or misshapen cells. In a few mildly deformed sh*lgl* embryos, disorganization appeared in restricted regions of the epithelium. However, the majority of sh*lgl* embryos manifested global epithelial disorganization with basal round cells and more ectodermal depressions compared to controls. Interestingly, similar ectodermal depressions were observed in sh*scribbled* embryos (Figure S2). Scribbled could function together with Lgl or be important for Lgl localization, explaining similar knockdown effects observed in fly epithelia ^20^. In contrast, reduced levels of apical *par3* and *par6* did not compromise ectodermal epithelia and no sh*lgl*-like deformities developed (Figure S3). In some severely damaged embryos, cells appeared to undergo lysis. However, these embryos also appeared in control groups (sh*gfp*), and we reasoned that this result may be due to the experimental procedure and not the specific gene knockdown. Indeed, the severely damaged embryos also appeared in similar numbers in all knockdown experiments, including those against *par3*, *par6*, *lgl* and *scribbled* polarity genes (Figures S2 and S3).

**Fig 2.**
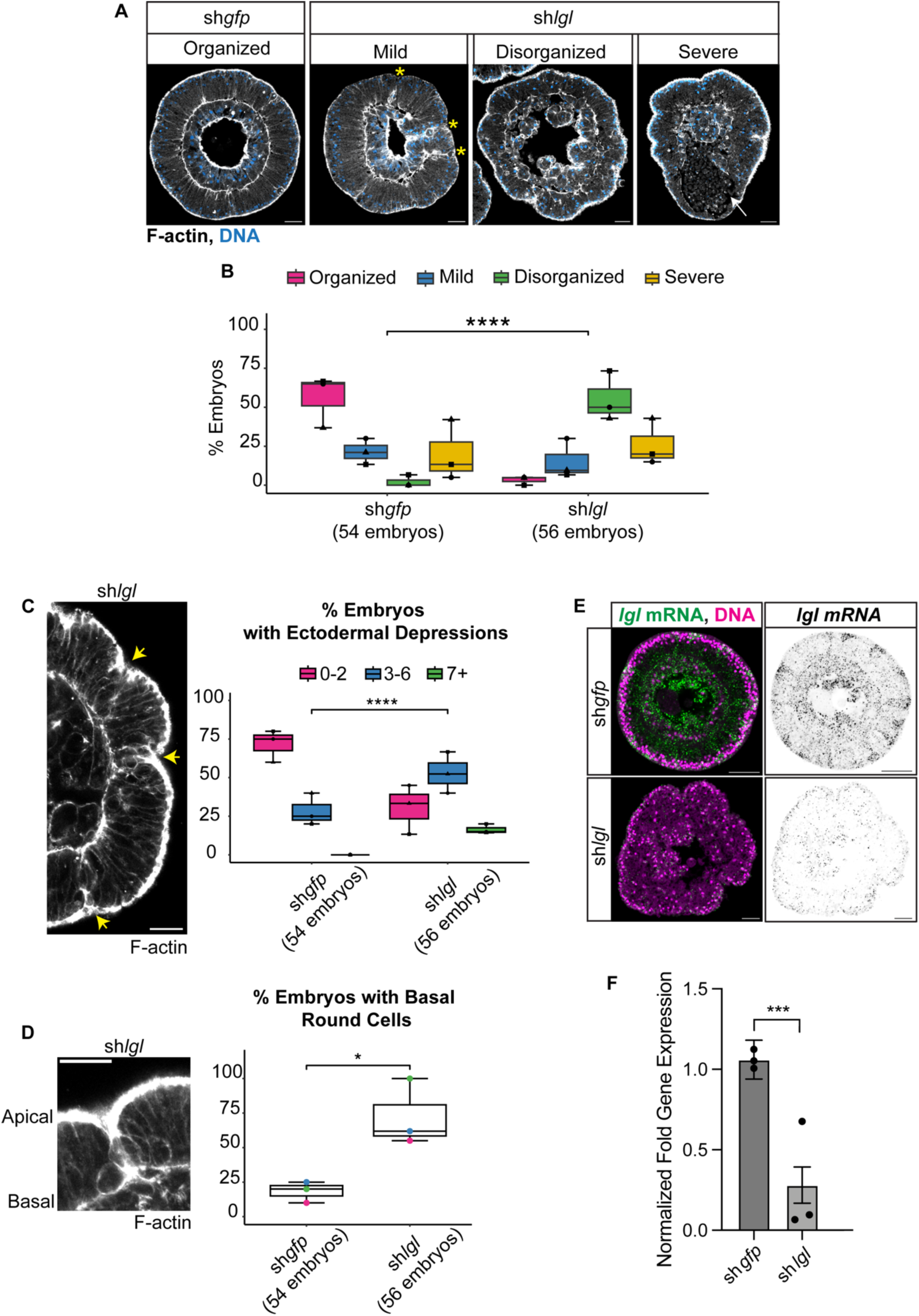
Expression of *lgl* is essential for epithelial organization in developing *Nematostella* embryos. (A) Ectodermal epithelia are disorganized in sh*lgl* embryos and manifest surface depressions not visible in the smooth epithelial layers of sh*gfp* control embryos. (B) Quantification of observed phenotypes. “Mild” refers to embryos with ectodermal epithelial disorganization in restricted areas of the ectoderm, shown near the yellow stars in “mild” embryo of (A). ”Disorganized” refers to embryos manifesting global epithelial disorganization with obvious depressions and misshapen cells throughout the ectoderm. “Severe” refers to embryos undergoing cell lysis pointed by the white arrow as an obliterated cell mass in “severe” embryo of (A). (Fisher’s exact test; ****p <0.0001). (C) Ectodermal depressions are more frequent in ‘disorganized’ sh*lgl* embryos and often correspond to areas with basal round cells pointed by the yellow arrows. (Fisher’s exact test; ****p <0.0001). (D) Basal round cells are prevalent in ‘disorganized’ sh*lgl* embryos. (ANOVA; *p =0.02). Symbols ▪▴● in (B) and (C) and colors blue, green and magenta in (D) correspond to embryo percentages from three independent experiments. (E-F) Expression of *lgl* is reduced in post-gastrula (28hpf) sh*lgl* embryos compared to sh*gfp* controls, as verified by *in situ* hybridization and qPCR of mRNA levels from embryos of three independent experiments. (Wilcoxon rank sum test; ***p< 0.001). Images show mid-sagittal sections from post-gastrula embryos. Scale bars: 20μm.

Taken together, our data suggest that expression of Lgl and Scribbled is essential for both epithelial organization of embryos and *de novo* epithelialization of cell aggregates.

## Epithelial disorganization in *lgl* mutants is not due to aberrant cell proliferation

We then sought to determine the mechanism by which knockdown of lateral polarity proteins disrupts epithelia. Specifically, we concentrated on examining the knockdown of *lgl* because of the robust disorganized phenotype seen in sh*lgl* embryos. Since its discovery as a tumor suppressor in the fly, Lgl has been implicated in both epithelial organization and cell proliferation ^19,20^. The *lgl* mutations transform the fly imaginal discs into lethal neoplasms and the optic primordia of the brain into neuroblastoma ^19,21^. Later work showed how Lgl together with Scribbled and Dlg, regulates cell polarity and epithelial organization in various fly epithelia ^20^. Similarly, knockout of one of the two Lgl homologues in mice (Lgl1) gives rise to disorganized neuroepithelial structures similar to human neuroectodermal tumors ^22^. Thus, we hypothesized that the epithelial disorganization observed in our *Nematostella* sh*lgl* knockdown embryos could be attributed to cell over proliferation. To examine whether cell proliferation control is disrupted, we fixed sh*gfp* control and sh*lgl* embryos at different stages during development and counted all stained nuclei. We also specifically identified mitotic nuclei using a phosphorylated Histone H3 (PH3) antibody and quantified the ratios of mitotic nuclei to total nuclei. Assuming there is one nucleus per cell, these quantifications reveal that both sh*gfp* and sh*lgl* embryos are composed of similar cell numbers throughout development (Figure 3 and Figure S4). These results further demonstrate that epithelial disorganization in *Nematostella* sh*lgl* mutants is not coupled to loss of cell proliferation control.

**Fig 3.**
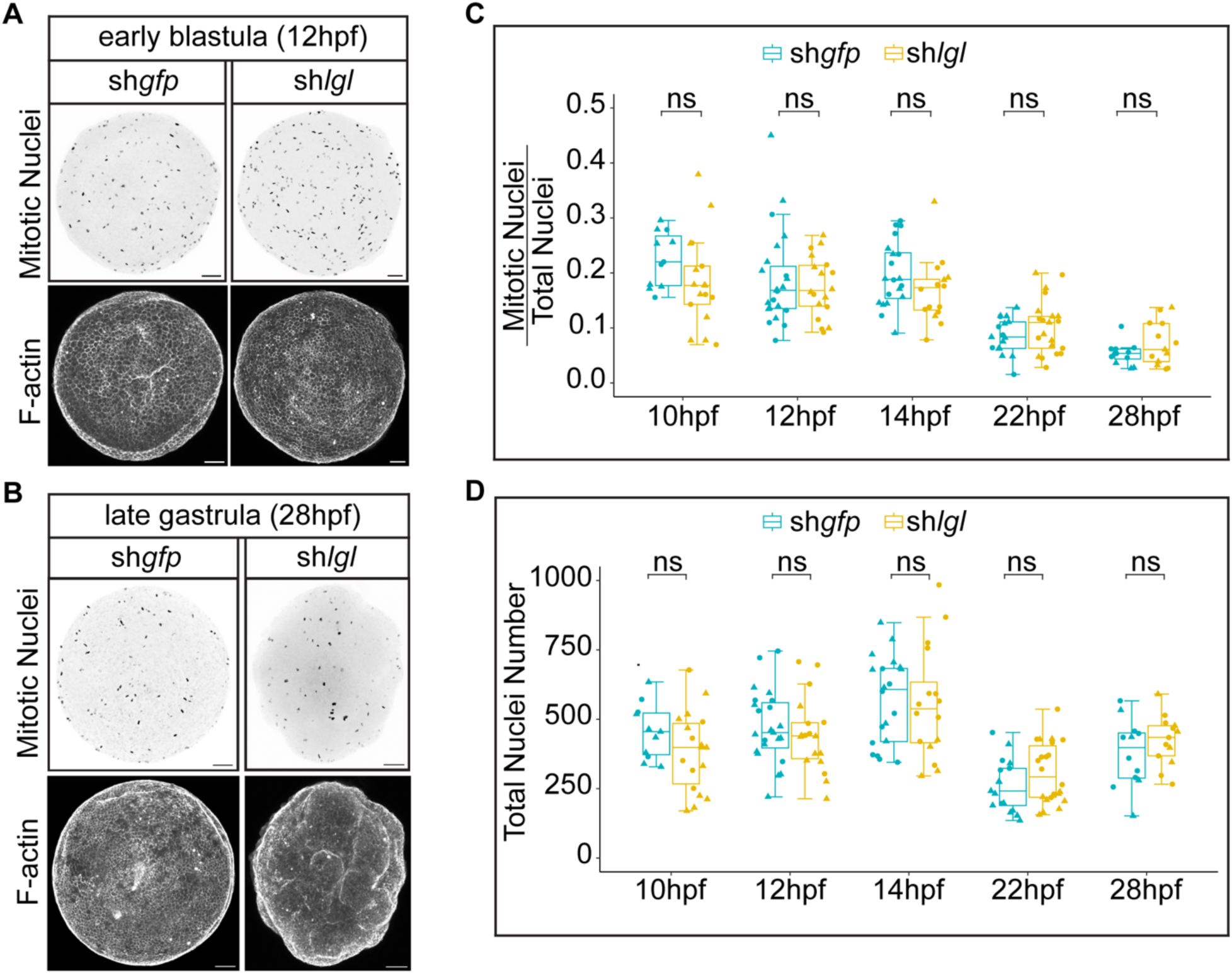
*lgl* mutant embryos do not exhibit cell over proliferation during development. (A) Mitotic nuclei stained with phosphorylated Histone H3 (PH3) antibody and corresponding F-actin-stained sh*gfp* control and sh*lgl* embryos at: (A) early blastula (12hpf) and (B) post-gastrula (28hpf) stages. (C) Quantification of ratios of mitotic nuclei/ total nuclei numbers in sh*gfp* control and sh*lgl* embryos across development. (D) Total nuclei quantification in sh*gfp* control and sh*lgl* embryos across development. Symbols correspond to average values from each embryo analyzed from two independent experiments. Number of sh*gfp* embryos analyzed per time point: 10hpf (11 embryos), 12hpf (23 embryos), 14hpf (20 embryos), 22hpf (17 embryos) and 28hpf (12 embryos). Number of sh*lgl* embryos analyzed per time point: 10hpf (17 embryos), 12hpf (18 embryos), 14hpf (16 embryos), 22hpf (21 embryos) and 28hpf (13 embryos). (Wilcoxon rank sum test; ns p> 0.05). Images show projections of whole embryos. Scale bars: 20μm. See also Figure S4.

### Basal domains are disorganized in *lgl* mutant epithelia

Lgl is enriched at the lateral cell contacts in *Nematostella* embryos ^15,16^. In epithelia of other species, Lgl restricts the localization of apical polarity proteins to the apical domain ^2,3^. Reduced Lgl levels are likely to disrupt epithelia by impairing cell polarity. To examine cell polarity in *Nematostella* embryos, we checked the localization of conserved proteins known to localize at the apical and basal domains in epithelial cells. We first examined whether apical domains are properly patterned in *lgl* mutant epithelia by checking the localization of Par3-GFP and Par6-GFP. In both sh*mcherry* controls and sh*lgl* embryos, Par6-GFP was enriched at the apical cortices and apical cell junctions (Figure 4A). Similarly, Par3-GFP was found at the apical cell junctions of both control and *lgl* mutants (Figure S5A). However, in contrast to controls, some cells in *lgl* mutants had Par6-GFP and Par3-GFP enriched at their basal domains as well (Figure 4A and Figure S5).

**Fig 4.**
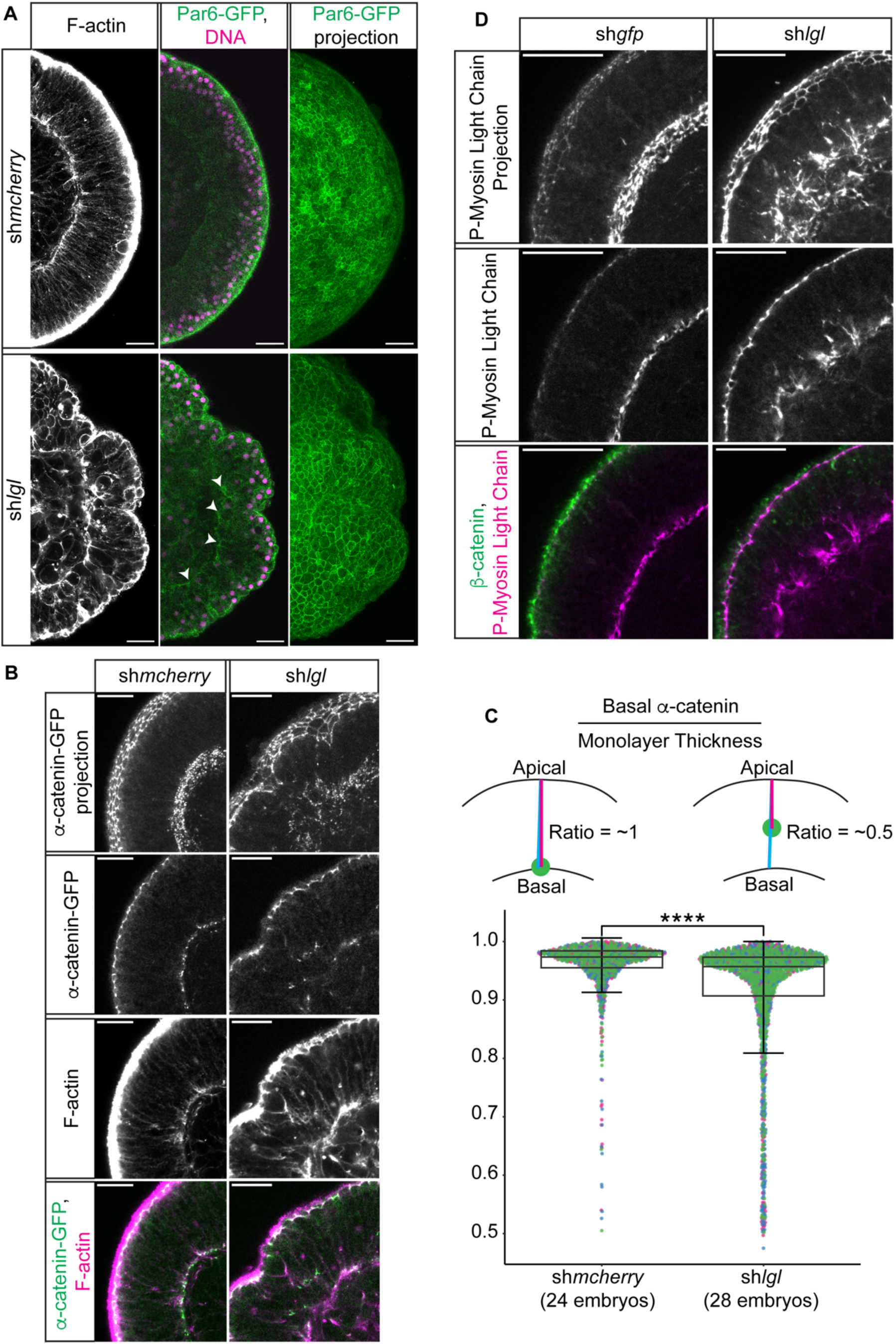
Basal polarity is impaired in *lgl* mutant epithelia. (A) Par6-GFP is enriched exclusively at the apical cell cortices and junctions, as shown in the mid-sagittal section and surface projection views of sh*mcherry* control embryos. In sh*lgl* embryos, Par6-GFP is enriched at apical cell cortices and junctions, similar to controls, as well as some basal regions of the ectodermal epithelium pointed by the white arrowheads. (B) α-catenin-GFP is enriched at both apical and basal junctions of the epithelium. In control sh*mcherry* embryos, apical and basal α-catenin-GFP mark the apical and basal domains of the monolayer at a constant distance along its thickness. In sh*lgl* embryos, basal α-catenin-GFP is enriched at variable distances from the apical domain along the monolayer. Projection is the sum of 15μm above and below the mid-sagittal section. (C) Quantification of basal α-catenin-GFP relative to the monolayer thickness in sh*mcherry* control and sh*lgl* embryos. Monolayer thickness is determined by F-actin which labels cell margins. Ratio = 1 indicates colocalization of basal α-catenin-GFP with the basal domain of the monolayer. Ratio < 1 indicates that basal α-catenin-GFP enrichment does not align along the basal domain and appears within a shorter distance from the apical domain. (Wilcoxon rank sum test; ****p< 0.0001). (D) Phosphorylated myosin light chain is enriched at apical and basal domains of ectodermal epithelia. In sh*gpf* control embryos phosphorylated myosin marks the basal domain of the monolayer in a consistent manner along its thickness, similar to the localization of basal α-catenin-GFP. In sh*lgl* embryos, basal enrichment of phospho-myosin is disorganized and appears at variable distances from the apical domain of the monolayer. Projection is the sum of 8μm above and below the mid-sagittal section. All images show mid-sagittal sections of post-gastrula embryos unless otherwise stated. Scale bars: 20 μm. See also Par3-GFP localization in Figure S5.

Enrichment of apical proteins in basal epithelial regions suggests a basal polarization defect. We further examined the integrity of basal domains by visualizing the localization of a GFP fusion to the junction protein α-catenin. *Nematostella* ectodermal epithelia are unique in that junctional Cadherin3 is enriched in both apical and basal domains ^18^. Similar to Cacherin3 localization, we found α-catenin-GFP enriched at both apical and basal cell junctions of wild type epithelia (Figure 4B).

However, in *lgl* mutants, α-catenin-GFP was not consistently found in the basal domain of the ectodermal monolayer, and was instead distributed at shorter distances from the apical domain (Figure 4B-C). This could be due to basal domains established at shorter cell lengths along the *lgl* mutant epithelium. Alternatively, α-catenin-GFP could be mis-localized along the cell’s lateral sides instead of a defined basal domain. In either case, mis-localization of α-catenin-GFP points to disorganized basal junctions in *lgl* mutants. Indeed, phosphorylated myosin that is typically enriched at defined junctions along the basal domain of wild type epithelia, was also found to be disorganized and distributed at various lengths along the *lgl* mutant monolayer (Figure 4D).

Taken together, examination of polarity and junction protein localization demonstrates that basal domains are disorganized in *lgl* mutant epithelia. The basal junctional markers, α-catenin-GFP and phosphorylated myosin, localize in shorter distance from the apical domain. These mispositioned basal junctions, if still functional, likely alter the contractile properties and behaviors of both individual cells and the ectodermal epithelium at large.

### Lgl is required for planar spindle orientation during early blastula cleavages

Disorganized actomyosin contractility and loss of cortical polarity could explain the appearance of misshapen cells and epithelial depressions in *lgl* mutant cells. A clear manifestation of disruption in actomyosin and polarity in the epithelium is misoriented cell divisions ^23,24^. In epithelia, mitotic spindle orientation is of paramount importance ^25–27^. To maintain the monolayer structure, spindles must be positioned parallel to the epithelial plane. To examine whether *lgl* knockdown affects spindle orientation in *Nematostella* epithelia, we examined the angle of mitotic spindles relative to the monolayer plane at various stages during development (Figure 5). Throughout development, we detected high variability of spindle position during prophase in both control and *lgl* mutant embryos. However, by metaphase, spindles in control embryos appeared to consistently align parallel to the epithelial plane. We did not detect any obvious differences between control (sh*gfp*) and sh*lgl* spindle orientation during early cleavages (Figure 5A, 6hpf). However, at the blastula stage, when cells are already organized into an epithelial monolayer, *lgl* knockdown significantly impaired spindle orientation (Figure 5B, 12hpf). After this stage, *lgl* mutant epithelia began to manifest deformities in their structure, and spindle orientation examination was restricted to regions where the epithelial plane was identifiable. In well-structured regions, spindles were parallel to the plane, even during later developmental stages (Figure 5C, 25hpf). This could be attributed to *lgl* knockdown penetrance effects, where Lgl protein levels are at different functional concentrations in across cells. Although knockdown of *lgl* mRNA was efficient throughout the embryo, maternally provided protein may not be depleted in all cells at the same time ^28^.

**Fig 5.**
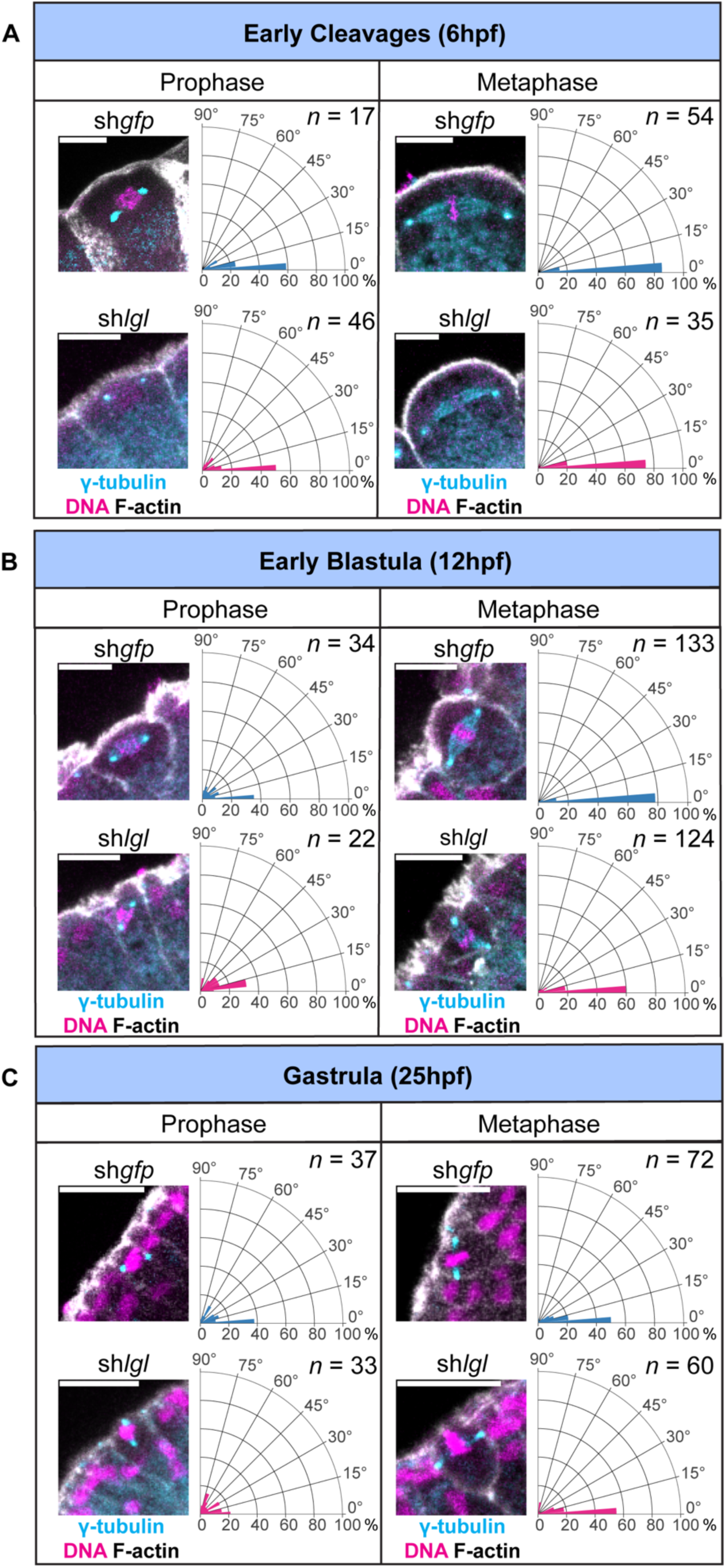
Lgl is required for planar spindle orientation during early blastula cleavages. (A-C) Prophase and metaphase spindles were identified by γ-tubulin enrichment at the spindle centrosomes in sh*gfp* control and sh*lgl* embryos during development. Corresponding fan plots show the percentage of spindles that are parallel (angle: 0 degrees) or perpendicular to the plane of the ectodermal epithelium (angle: 90 degrees). Prophase spindles show a randomized orientation relative to the epithelial plane throughout development in both sh*gfp* and sh*lgl* embryos. Metaphase spindles align with the plane of the epithelium in sh*gfp* embryos throughout development. In sh*lgl* embryos, a significant percentage of spindles deviates from the epithelial plane during early blastula cleavages (12hpf) shown in (B). (Wilcoxon rank sum test; p= 0.009). (C) By gastrula stage, analysis of sh*lgl* spindles was restricted to well-structured embryo regions with identifiable epithelial planes. In well-structured regions, sh*lgl* spindles were found oriented parallel to the epithelium. *n* indicates the number of spindles. Number of sh*gfp* embryos analyzed per time point: 6hpf (10 embryos), 12hpf (25 embryos) and 25hpf (20 embryos). Number of sh*lgl* embryos analyzed per time point: 6hpf (12 embryos), 12hpf (21 embryos) and 25hpf (17 embryos). With the exception of 12hpf metaphase spindles, the median angle of spindle orientation is not significantly different between sh*gpf* and sh*lgl* embryos (Wilcoxon rank sum test; p> 0.05). Scale bars: 10μm.

In conclusion, we identified spindle orientation defects in cells of early blastula *lgl* mutant embryos. These defects could drive cells to undergo oblique or orthogonal divisions relative to the epithelium such that one daughter cell positions its apical “domain” towards the basal domain of the monolayer. Interestingly, we did not identify any Par3-GFP or Par6-GFP enrichment within the ectodermal monolayer. Assuming that daughter cells appear in mirror symmetry relative to each other after orthogonal divisions, the plane of cleavage could explain the appearance of Par3-GFP and Par6-GFP apical proteins in the basal domain of the ectodermal epithelium and basal proteins like α-catenin-GFP and phosphorylated myosin within the epithelium (Figure 4, S5). Mispositioned mitotic spindles could lead to the apicobasal polarity defects observed within some regions of the ectodermal epithelium.

### Mispositioning of the basal body in *lgl* mutant epithelia

An unexpected observation in *lgl* mutant embryos was the mispositioning of ciliary basal bodies. During our examination of epithelial polarity, we serendipitously discovered that a commercial antibody previously used for β-catenin labelling marked both the apical cell junctions and the basal body of *Nematostella* ectodermal epithelia (Figure S6). By late blastula stage (18hpf), embryos develop motile cilia and basal bodies become evident at the apical domain of their ectodermal epithelia. Closer inspection revealed that β-catenin localizes in a ring-like pattern around the basal body marked by y-tubulin near the center of the cell’s apical region. Interestingly, in sh*lgl* mutants, the basal body was found mispositioned in close proximity to the lateral plasma membrane (Figure 6A-B). We quantified the position of the basal body relative to the nearest cell contact in cells from sh*gfp* controls and sh*lgl* mutants by using both β-catenin and y-tubulin markers. We found a significantly higher distribution of basal bodies near the cell contact in sh*lgl* mutant epithelia compared to controls (Figure 6C-D). Considering that the basal body is an apically-docked cell organelle and apical polarity appeared normal in *lgl* mutant epithelia, its mispositioning is particularly intriguing.

**Fig 6.**
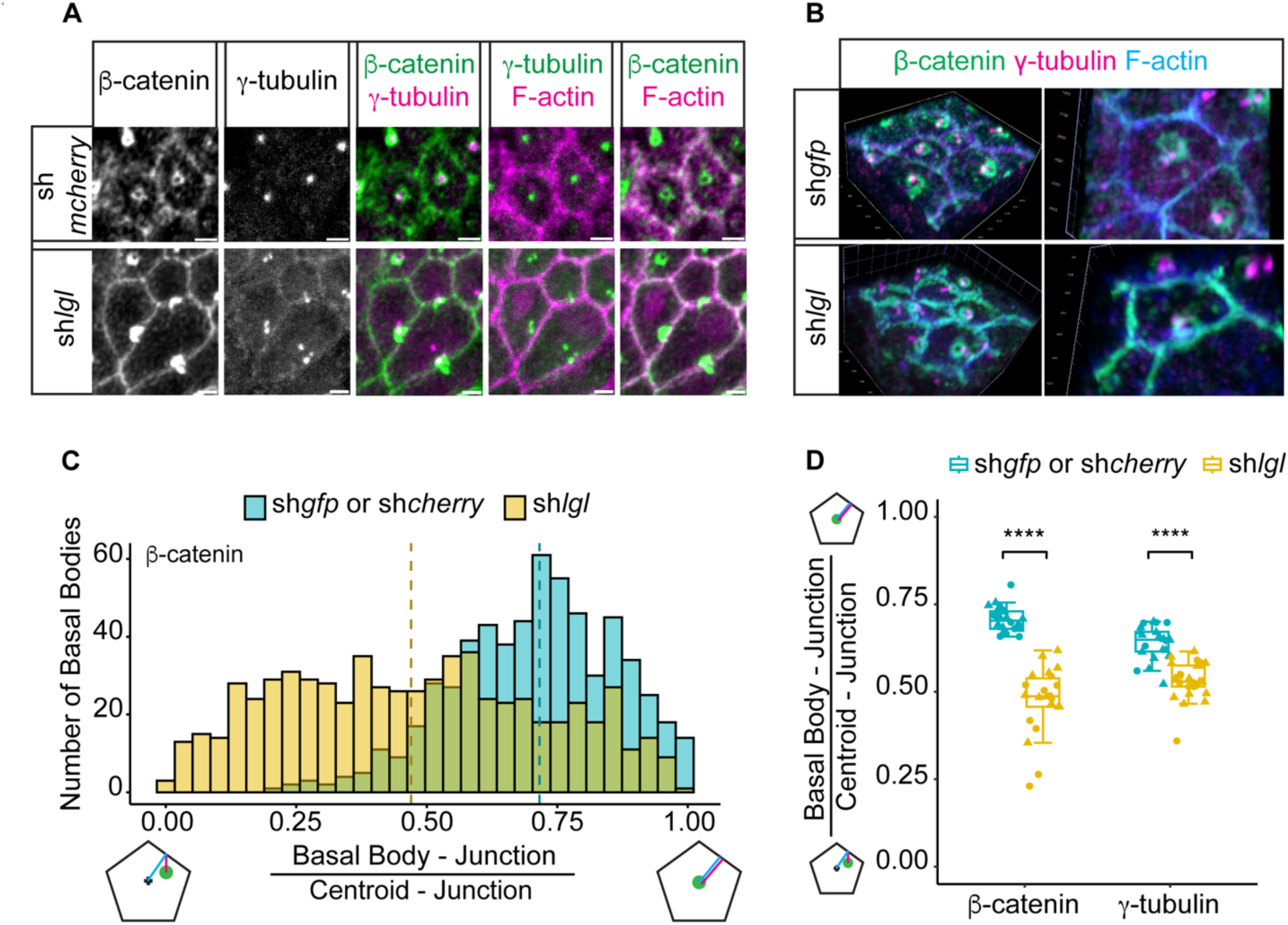
The basal body is mispositioned in the ectodermal epithelia of *lgl* mutant embryos. (A) Position of the basal body in sh*gfp* control and sh*lgl* embryos as indicated by the enrichment of β-catenin and γ-tubulin relative to the junctions of ectodermal epithelial cells. (B) Closeup 3D projection views of β-catenin enrichment near the basal body as indicated by enrichment of γ-tubulin in sh*gfp* and sh*lgl* embryos. Basal bodies are near-center-localized in sh*gfp* controls and closer to the cell junctions in sh*lgl* embryos. (C) Distribution of basal body positions relative to the cell center and cell junction in sh*gfp* control and sh*lgl* embryos. Medians are: 0.716 for sh*gfp* and 0.47 for sh*lgl* basal bodies. Analysis was based on analysis of β-catenin-marked basal bodies and cell junctions. (D) Average ratios of β-catenin and γ-tubulin-marked basal body positions relative to the cell center and cell junction in sh*gfp* control and sh*lgl* embryos. Symbols ▴● correspond to ratios from two independent experiments. Ratio = 1 indicates basal body position closer to the center of the cell. Ratio < 0.5 indicates basal body position closer to the cell junction. Number of β-catenin-marked-basal bodies analyzed in 20 sh*gfp* embryos: 601. Number of β-catenin-marked-basal bodies analyzed in 22 sh*lgl* embryos: 661. Number of γ-tubulin-marked-basal bodies analyzed in 22 sh*gfp* embryos: 660. Number of γ-tubulin-marked-basal bodies analyzed in 26 sh*lgl* embryos: 780. (Wilcoxon rank sum test; ****p<0.0001). Images and analysis are from post-gastrula embryos. Scale bars: 2μm.

## Discussion

Epithelia are the first organized tissues to appear during animal development ^1^. Epithelia emerge as cells divide, differentiate and migrate to form distinct territories and organs. *Nematostella* embryos self-organize into epithelial layers early during development and provide an excellent system for studying epithelial organization in isolation, independent of the confounding effects of signaling from neighboring organs and tissues ^14,15^. Additionally, dissociated *Nematostella* embryonic cells self-organize into a complete animal after aggregation ^8^. We reasoned that epithelialization of aggregates likely requires a similar set of conserved polarity proteins that are required to form epithelia in other animals. We tested four polarity proteins and found that aggregates from *Nematostella* cells with reduced *lgl* expression failed to form smooth epithelia and eventually disintegrated. This striking result first confirms that successful epithelialization needs to occur before an aggregate can further develop into a complete animal. Secondly, lateral polarization is essential for aggregate epithelialization. Knock down of two apical polarity proteins, Par3 and Par6, did not impair epithelialization and aggregates eventually developed into living tissues seven days post dissociation. Is lateral polarity only required for *de novo* epithelialization of aggregates? We found that *scribbled* and *lgl* knockdown also impaired embryonic epithelia. It is likely that embryos lacking expression of these two lateral proteins are structurally compromised and their dissociation gives rise to cells that lack the potential to epithelialize. A future experiment would be to trigger expression of lateral polarity proteins in dissociated cells from mutants post-aggregation and examine whether their presence restores epithelialization. Interestingly, knockdown of Par3 and Par6 proteins did not impair either embryonic or aggregate epithelialization. Additionally, GFP fusions to Par3 and Par6 localized normally to apical domains in *lgl* mutant embryos, indicating normal apical polarity even when epithelia were highly disorganized. This result suggests that intact apical polarity may not be sufficient to override basolateral polarity defects and maintain epithelial integrity. Conversely, loss of lateral polarity is detrimental for both embryonic and aggregate epithelialization.

Indeed, embryos with reduced *lgl* expression develop surface depressions that accompany basal polarity defects. Specifically, we found three features that destabilize the basal domain of the ectodermal monolayer. First, round non-mitotic cells occupy the basal domain. This is in contrast to normal epithelia, where mitotic cells undergo cortical rounding in the apical domain and no round cells appear in the basal domain ^25^.

Second, some cells develop apical character in the basal region of the ectodermal epithelium. Par6 and Par3 enrichment is never encountered in the basal domain of normal epithelia. Studies in other systems have uncovered a robust mechanism that blocks apical proteins from localizing to the basal domain of epithelial cells ^2,3^. Although not yet examined in *Nematostella*, a similar mechanism of apical protein exclusion from the basal domain may also exist. Finally, basal junctions were not confined in the basal domain of the monolayer. Instead, we found both α-catenin-GFP and phosphorylated myosin localized at various lengths from the apical domain. This could be due to mis-localization of basal junctional proteins along normally elongated epithelial cells. Alternatively, the cells develop basal polarity but are shorter in length. Together, these three features explain the formation of a bi-layered structure composed of basal cells with reversed polarity to that of the apical cells stacked above them.

A mechanism that could give rise to basal cells with reverse polarity is misoriented cell division. When mitotic spindles are positioned in oblique or orthogonal angles relative to the epithelial plane, one of the daughter cells ends up in the basal domain of the monolayer. If one daughter cell polarizes in mirror symmetry to the other, we would expect to find cells with their apical proteins in the basal domain and their basal proteins buried within the monolayer. We did find a higher number of misoriented cleavages in *lgl* mutants during early blastula. However, we also detected normal cleavages during development, possibly due to sufficient Lgl protein levels in some cells. Nevertheless, few misoriented cell divisions early during development could strongly impact epithelial organization.

We were surprised to find that although the apical domains of *lgl* mutant epithelia were properly polarized, basal bodies of their cilia were mispositioned. Depending on the phase of the cell cycle, centrioles form spindle poles during mitosis and basal bodies during interphase ^29^. In normal epithelia, basal bodies are anchored within close range to the centroid of the apical domain. In *lgl* mutants, we observed a high proportion of basal bodies positioned very close to the lateral plasma membrane. Disruption of cytoskeletal organization is likely to impair basal body migration or docking to the apical cell membrane. A similar defect in basal body anchoring to the lateral plasma membrane was observed when quail oviducts were treated with Benzodiazepines. These drug treatments also impaired cortical actomyosin and vesicle trafficking, suggesting that basal body mispositioning could be due to a migration defect ^30^. Misorientation of the mitotic spindle during early blastula cell divisions and mispositioning of the basal body in interphase cells of *lgl* mutants both point to a potential centriole movement defect.

It is unclear whether Lgl has a direct role on centriole positioning in *Nematostella* epithelia. In human embryonic kidney cells, Lgl2 binds and regulates localization of Pins (Partner of Inscutable), an LGN homologue which has a role in spindle pole orientation^31^. Lgl is also implicated in the regulation of symmetric cell divisions in fly epithelia ^32,33^. However, in fly, it is cortical release of Lgl that drives proper cell division. Retention of Lgl on the cortex impairs spindle positioning. Lgl cortical release frees Dlg to interact with Pins/LGN in the lateral domain and promote planar orientation of the spindle.

Moreover, *lgl* mutant follicular epithelial cells and syncytial embryos assemble normal spindles and undergo normal chromosome segregation respectively, suggesting that in the fly Lgl does not have a direct role in chromosome segregation ^32^. Similarly, Lgl may not play a direct role in spindle and centriole positioning in *Nematostella*. Other factors, such as well-organized cortical actomyosin is critical for planar spindle orientation in epithelial tissues and could also play a role in *Nematostella* embryos.

We propose that in *Nematostella*, the lateral polarity protein Lgl is important for epithelialization mainly through its role in organizing the cytoskeleton. Cell shape change, misorientation of the mitotic spindle, and mispositioning of the ciliary basal body in *lgl* mutants could be all attributed to a failure of cytoskeletal organization. Early studies in *Drosophila* showed that Lgl binds to and regulates the localization of myosin II ^34,35^. Cells in *lgl* mutant embryos and imaginal discs are cuboidal or round instead of long and columnar and as a result fail to establish extended lateral contacts. It was proposed that the Lgl-myosin complex stabilizes newly formed lateral cell contacts and maintains cell shape ^34,36^. Human Lgl also binds to myosin II, and in *lgl* knockout mice brains the organization of actin and localization of myosin II are perturbed ^22,37^.

Interestingly, we did not find any correlation between Lgl and cell proliferation control in early *Nematostella* embryos. Since its discovery as a tumor suppressor in the fly, Lgl has been shown to be important for control of cell proliferation in various organs including imaginal discs, brain, eye and egg chamber ^19,20,38–40^. In mammals, Lgl affects cell numbers in the brains of mouse embryos and human mammary epithelial cells grown in a 3-dimensional extracellular matrix ^22,41^. However, in some of these cases, cell division is coupled to tissue differentiation. For example, knockout of one of the two Lgl homologues in mice (Lgl1) leads to the loss of asymmetric inheritance of the Notch signaling antagonist Numb, and neural progenitors fail to differentiate and continue to proliferate into tumor-like structures ^22^. It will be important to examine the effect of *lgl* expression later during *Nematostella* development, when cells are undergoing differentiation and organ development.

Epithelialization is a fundamental tissue organization process that occurs in all developing animals. However, it is context dependent with variations across animals, tissues within animals and stages of development ^1,2^. Here, we show that lateral polarity is essential for organization of epithelial layers during both embryonic development and aggregate organization from dissociated cells. Although, many factors are at play during epithelialization including adhesion, we have found an important role for lateral polarization. In the absence of defined lateral polarity, adhesion proteins are not sufficient to organize cells into an epithelium. Similarly, cysts developing in suspension from Madin Darby Canine Kidney (MDCK) cells require formation of both lateral cell contacts and basal lamina during *in vitro* epithelialization ^42^. Establishment of lateral polarity may be a central feature of self-organization in other epithelializing systems.

## Acknowledgments

We are grateful to Dr Lampros Panagis (Amherst Biology Imaging Center) and Phillip Zhou for training and help with imaging, Dr Nicholas Horton and Clara Page for advice on statistical analysis, Dr Cheng-Yi Chen for qPCR advice, and Caroline Towse and the Biology Department of Amherst College for animal maintenance and administrative assistance. Our work is supported by Amherst College, an NSF-MRI grant 2117798 and a NIH grant R15GM141979-01.

## Materials and Methods

## Animal and embryo handling

*Nematostella vectensis* animals were cultured in the dark in 12 parts per thousand (ppt) artificial sea water (SW) (Sea Salt: Instant Ocean) at 17°C, were fed weekly with freshly hatched Artemia salina brine shrimp larvae and were spawned every 3 weeks as previously described ^43^. Eggs were de-jellied soon after spawning and were either injected with a mix of mRNA and shRNA or electroporated with shRNA, as described below ^15,44^. Treated eggs were fertilized immediately after injection or electroporation and embryos were kept at room temperature (RT). The transgenic line *mat>Par6-GFP* provided embryos for the Par6-GFP localization studies and was previously described^15^.

### Gene knockdown by shRNAs

shRNAs were designed and prepared as previously described ^44^. Sequences of genes of interest were submitted to siRNA Wizard web interface (InvivoGen; http://www.invivogen.com/sirnawizard/design.php) with a 19 nucleotide-long motif size. Sequences of approximately 50% GC content were checked by BLASTn against the *Nematostella vectensis* genome and after confirming specificity were used for primer design and ordering from Integrated DNA Technologies (see full list of shRNA primer sequences in Table S1). shRNAs against *gfp* and *mcherry* were used as negative controls. Forward and reverse primers were annealed after heating to 95°C for 5min and allowing the mix to cool at 22°C. The annealed DNA template containing the T7 promoter and shRNA sequence (included in the primer design) was then used for *in vitro* shRNA transcription via the Ampliscribe T7-Flash Transcription Kit. shRNAs were purified using the Direct-zol RNA MiniPrep Kit (Zymo Research), were eluted in molecular biology-grade water and were quantified by a Nano Drop spectrophotometer. shRNAs (500ng/μl in 15% Ficoll PM400/SW) were electroporated into dejellied eggs as previously described ^44^.

### Quantification of gene expression after shRNA knockdown

Knockdown of gene expression was confirmed by qPCR on three biological replicates. At 28hpf, 100 embryos were collected from each treatment, spun down to a pellet with SW removed, rapidly frozen in liquid Nitrogen and stored at -80°C. Total RNA was extracted and purified with the Direct-zol RNA Miniprep Kit (Zymo Research). Total cDNA was prepared from the purified RNA using the iScript cDNA Synthesis Kit (Bio-Rad). Quantification of expression was done by qPCR that was carried out in three technical replicates for each sample, using the SsoAdvanced Universal SYBR Green Supermix (Bio-Rad) and the qPCR primers listed in Table S1 on a CFX Duet Real-Time PCR System (Bio-Rad). ATP synthase was chosen as the endogenous reference gene for normalization due to its invariability in expression across treatments. Relative changes in gene expression were calculated with the ΔΔCq method, and statistical comparisons between control (sh*gfp*) and polarity mutants were done using the Wilcoxon rank sum test.

Embryos were also examined for *lgl* knockdown by Fluorescent *In Situ* Hybridization (FISH) as previously described ^45^. Briefly, embryos were collected at 28hpf and were fixed at RT in ice-cold 4% paraformaldehyde and 2.5% glutaraldehyde in SW for 90s, followed by fixation in 4% paraformaldehyde in SW for 1h at RT. After washing in PBS/0.1% Tween20, embryos were dehydrated through a PBS/Methanol gradient and stored at -20°C. Embryos were washed in 3% H_2_O_2_ in 90% Methanol for 1h at RT under light and rehydrated through a Methanol/PBS gradient. After Proteinase K treatment (80μg/ml for 3min) embryos were hybridized with 1ng/μl DIG-labeled *lgl* RNA probe at 60°C for 48h followed by overnight incubation with pre-absorbed sheep anti-DIG-POD Fab (Roche, 1:1000) in 5% sheep serum (Sigma)/0.5% blocking reagent (Roche)/0.1%DMSO/TNT buffer at 4°C. Embryos were incubated with 1:2000 Draq5 (Cell Signaling 4084) in PBS/0.1% Tween20 to stain nuclei and were fluorescently labelled with a TSA Cyanine 3 (Cy3) detection kit (Akoya). Finally, embryos were dehydrated through an Isopropanol gradient and mounted in a 2:1 mixture of Benzyl Benzoate: Benzyl Alcohol. The *lgl* RNA probe (1000 nucleotide-long) was synthesized from cDNA (prepared from gastrula stage RNA) using primers listed in Table S1 and was labelled with Digoxigenin (DIG) using the T3 RNA Polymerase system (Promega).

### Embryo injection with mRNA and shRNA constructs

De-jellied eggs were injected with FITC fluorescent tracer, shRNA (500 ng/μl prepared as described above) and 600-1000 ng/μl mRNA that was *in vitro* transcribed (mMessage mMachine T3 kit, Invitrogen, AM1348) from constructs Par3-GFP and α−catenin-GFP described elsewhere ^15^. In control experiments, eggs were injected with shRNA targeting *mcherry*. De-jellied eggs were injected using pulled glass capillaries with the Eppendorf FemtoJet 4i system and were fertilized immediately after.

### Embryo dissociation and aggregation

Embryos were dissociated and their cells reaggregated as previously described ^8^. Briefly, 30 post-gastrula stage embryos (28hpf) were collected in 100μl SW and to that 200μl of Ca^2+^/Mg^2+^-free artificial sea water (27 g/L NaCl, 1 g/L Na_2_SO_4_, 0.8 g/L KCI, 0.18 g/L NaHCO_3_ in Milli-Q water) was added. Embryos were dissociated by pipetting 100 times through a p200 micropipette. The suspension was then passed through a 40μm porosity Flowmi Cell Strainer (Sigma) into a clean 2mL tube. The strained suspension volume was adjusted up to 2mL with SW and was centrifuged for 30min at 2.700 x *g* (Sorvall Legend Microcentrifuge 21). The pellet/aggregate was then moved into a petri dish containing SW and was cut into 4 equal sized pieces using a nose hair fixed to a glass Pasteur pipette. Individual aggregates were either left to grow in individual dishes filled with SW at RT, or imaged live, or fixed and stained for F-actin and DNA as described below.

### Fixation of embryos and immunohistochemistry

For all antibody staining procedures, *Nematostella* embryos were fixed as previously described ^18^. Briefly, embryos were fixed with 4% paraformaldehyde/PBS for 1h at 4°C, followed by incubation on ice with ice-cold acetone for 7min. After seven washes in PBS/0.2% TritonX-100, embryos were incubated in blocking solution (PBS/0.2% TritonX-100, 20% normal goat serum (Sigma), 1% bovine serum albumin (BSA), 1% DMSO) for 2h at RT before overnight incubation in the same buffer with the chosen primary antibodies at 4°C. Primary antibodies were used in the following concentrations: mouse phospho-Histone H3 antibody (1:1000, Sigma 05-806), mouse α-tubulin antibody (1:1000, Sigma T6074), mouse γ-tubulin antibody (1:1000, Sigma T6557), rabbit β-catenin antibody (1:200; Sigma C2206) ^18,46^and mouse phospho-myosin light chain 2 antibody (1:200, Cell Signaling 3671). After six washes in PBS/0.2% TritonX-100, embryos were incubated with blocking solution for 2h at RT, followed by overnight incubation in blocking solution with the chosen secondary antibodies at 4°C. The following secondary antibodies were all used in 1:1000 concentration: goat anti mouse Alexa Fluor 488 antibody (Invitrogen, A11001), goat anti-mouse Alexa Fluor 568 antibody (Invitrogen, A11004), goat anti-rabbit Alexa Fluor 488 (Invitrogen, A11008) and goat anti-rabbit Alexa Fluor 555 (Invitrogen, A21428). DNA was stained with either DAPI (1μg/ml, Sigma D9542) or Draq5 (1:2000, Cell Signaling 4084) and F-actin was stained with Alexa Fluor 488, 555 or 647 phalloidin (1:200, Invitrogen A12379, A34055 or A22287, respectively) that were added in the overnight secondary antibody incubation at 4°C. After two washes in PBS/0.2% TritonX-100, embryos were dehydrated in an Isopropanol gradient and mounted in 2:1 mixture of Benzyl Benzoate: Benzyl Alcohol.

For F-actin and DNA staining of embryos, fixation was carried out as previously described ^47^. Aggregates were fixed with 4% paraformaldehyde/PBS by overnight incubation at 4°C. Staining of DNA and F-actin and mounting of embryos and aggregates was performed as described above.

Embryos injected with GFP-expressing constructs were fixed on ice in 4% paraformaldehyde and 0.1% glutaraldehyde in SW for 30min. Embryos were then washed three times with TBS/0.2% Triton-X100 and two times with TBS/1% BSA for 5min each, before overnight incubation with TBS/1% BSA supplemented with DAPI (1μg/ml) and Alexa Fluor 555 phalloidin (1:500) at 4°C. Just before imaging, embryos were washed twice with TBS/02% Triton-X100 and once with TBS for 10min each, were then taken though a Glycerol/TBS gradient, and were mounted in 90% Glycerol.

### Imaging

Fixed embryos were imaged with a 40x oil objective (1.30NA) using an inverted confocal Nikon Ti Eclipse microscope, and with a 40x water objective (1.20NA) on an inverted Zeiss LSM980 confocal microscope. For the cell proliferation analysis nuclei were imaged with a 20x objective (0.8NA), and airyscan images of the basal bodies were obtained with an oil 63x High Strehl ratio objective (1.4NA), both using the Zeiss LSM 980 microscope.

For time lapse imaging of live aggregates, samples were placed in glass-bottom MatTek dishes containing SW, and lids custom-made with 1.5 thickness cover slip (constructed by L. Panagis). Aggregates were imaged with differential interference contrast (DIC) using a 10x objective (0.45NA) on an inverted widefield Axio Observer 7 Zeiss microscope equipped with a Hamamatsu Orca Flash 4.0 camera. Images were collected at RT with a 30min interval and an adjusted Z-stack to include the whole aggregate.

### Image analysis

Images were processed using Fiji (ImageJ). Depending on the experiment, images were analyzed as follows. For mitotic nuclei/total nuclei analysis, whole embryos were imaged with a Z step of 2μm between slices. Noise and background were subtracted. In areas with minimal nuclei overlap, Z-series were displayed as a stack of maximum Z-projections of 2-3 slices and the stack was used for analysis; in regions with significantly high number of nuclei and risk of overlap, single slices were analyzed. In each slice or stack, total nuclei (DAPI or Draq5) were counted using the cell counting Bio App of the Zen image processing software selecting for area (2-40 μm^2^) and circularity (0-1.0). In the same slice or stack, mitotic nuclei (phospho-Histone H3) were counted manually.

For α-catenin-GFP localization analysis, a slice from the middle of post-gastrula embryos was selected (mid-sagittal). Analysis was performed on regions with no ectodermal depressions. Using as reference the image where cell boundaries are visible by F-actin staining, a 250 pixels x 250 pixels square region of interest was selected and cropped. Four separate regions were randomly selected for each embryo and all α-catenin-GFP points of enrichment were analyzed. Apical and basal domains of the ectodermal monolayer were outlined and interpolated into 0.05 pixel intervals. All visible points of α-catenin-GFP enrichment were identified and the shortest distance between α-catenin-GFP and the apical domain of the embryo was measured. A ratio of ((α-catenin) - (apical domain)) distance / monolayer thickness at that region was calculated. A ratio of ∼1 indicates basal α-catenin-GFP enrichment and a ratio of ∼0.5 indicates α-catenin-GFP enrichment halfway across the ectodermal monolayer.

For mitotic spindle orientation analysis, embryos were imaged with a Z step of 1μm between slices, and 2-4 slices of the Z-stack were displayed as maximum projections. To quantify orientation of the spindle, the angle between manually drawn lines was measured; one line was drawn parallel to the epithelial plane (identified by F-actin) and the other through the spindle poles (marked by γ-tubulin). Angle measurement values close to 0° signify near planar spindle orientation and values >0° and <90° represent oblique to near orthogonal spindle positioning.

For basal body localization analysis, post-gastrula embryos were imaged with a Z step of 0.5μm between slices, and 7-12 slices of the Z-stack were displayed as maximum projections. The basal body was marked using an oval Region Of Interest (ROI) tool and all cell junctions were outlined using a multipoint ROI tool. The centroid coordinates of both selected ROIs were obtained, and the cell junction outlines were interpolated into 0.05 pixel intervals. Then, the shortest distance between each centroid coordinate and a cell junction was measured. A ratio of (basal body) - (cell junction) distance / (centroid) – (cell junction) distance was calculated. A ratio of ∼1 indicates basal body localization at or near the cell centroid, and a ratio of ∼0 indicates basal body localization at or near the cell junction. In embryos with labelled β-catenin, basal body and cell junctions were visible in the same channel since β-catenin is enriched in both compartments. In embryos with labelled γ-tubulin, the basal body was marked by γ-tubulin and the cell junctions were marked by F-actin in a separate channel.

### Statistical analysis

Data analysis and plots were prepared using Prism 10 as well as R 4.3.1 with tidyverse, ggplot2, ggpubr, and plot_ly packages. Wilcoxon rank sum test, Fisher’s exact test, and ANOVA were used for testing of significance. Significance level, alpha, was set to 0.05.

---

**Fig S1.**
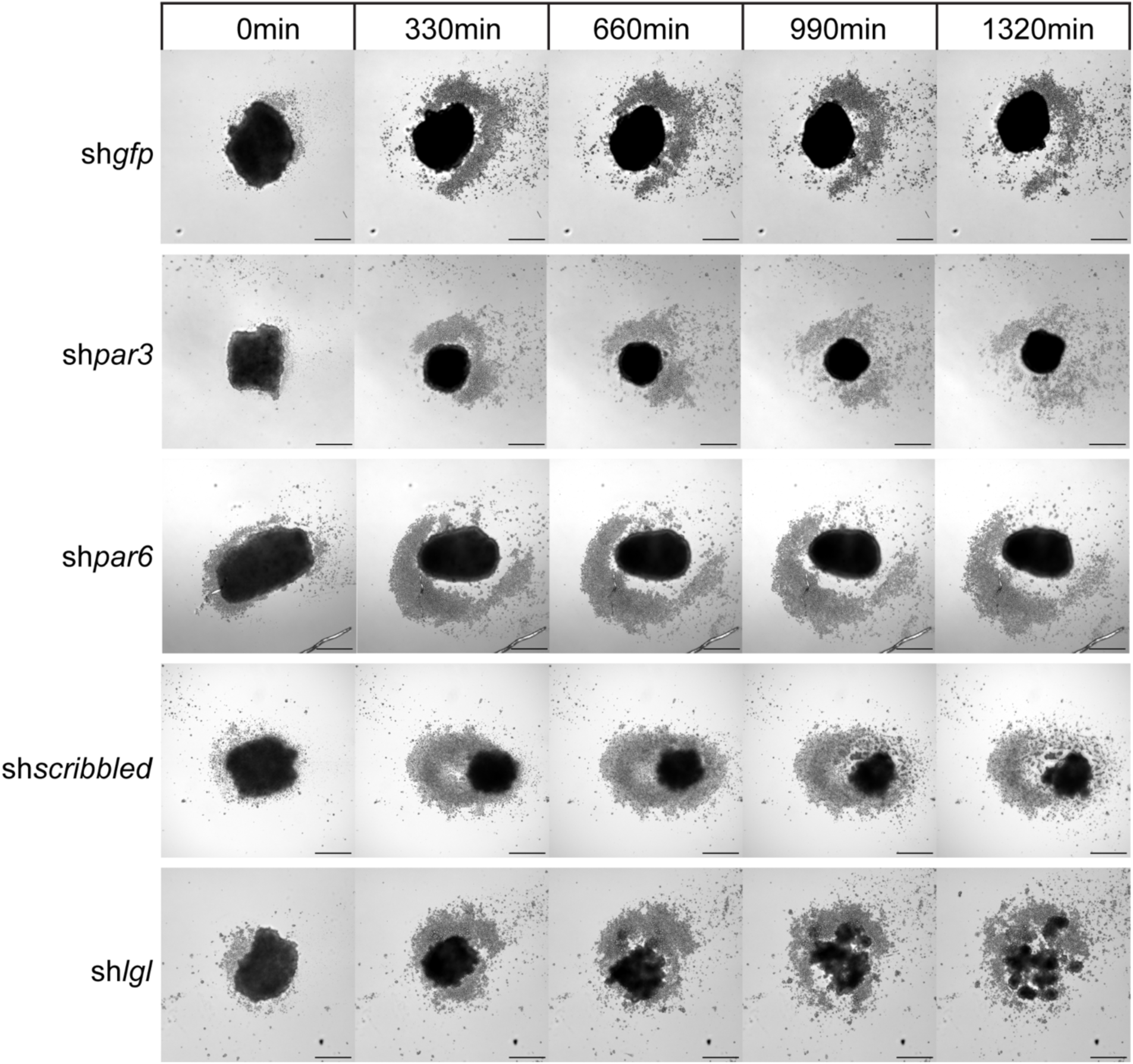
Lateral polarity proteins are essential for epithelialization of *Nematostella* embryonic aggregates. Still images from time lapse recording of single aggregates as they organize into epithelia in time. The majority of the aggregated cells from dissociated sh*gfp*, sh*par3*, sh*par6* and sh*scribbled* embryos compact into a smooth arrangement and stay together for the whole duration of the imaging (1320min or 22h post dissociation). In contrast, aggregates from dissociated sh*lgl* embryos begin to fall apart halfway through imaging (660min or 11h post dissociation). The sh*lgl* aggregate disintegrates into progressively smaller fragments that remain separate by the end of imaging. Images show projections of whole aggregates. Time lapse imaging of each aggregate was performed twice. Scale bars: 250μm.

**Fig S2.**
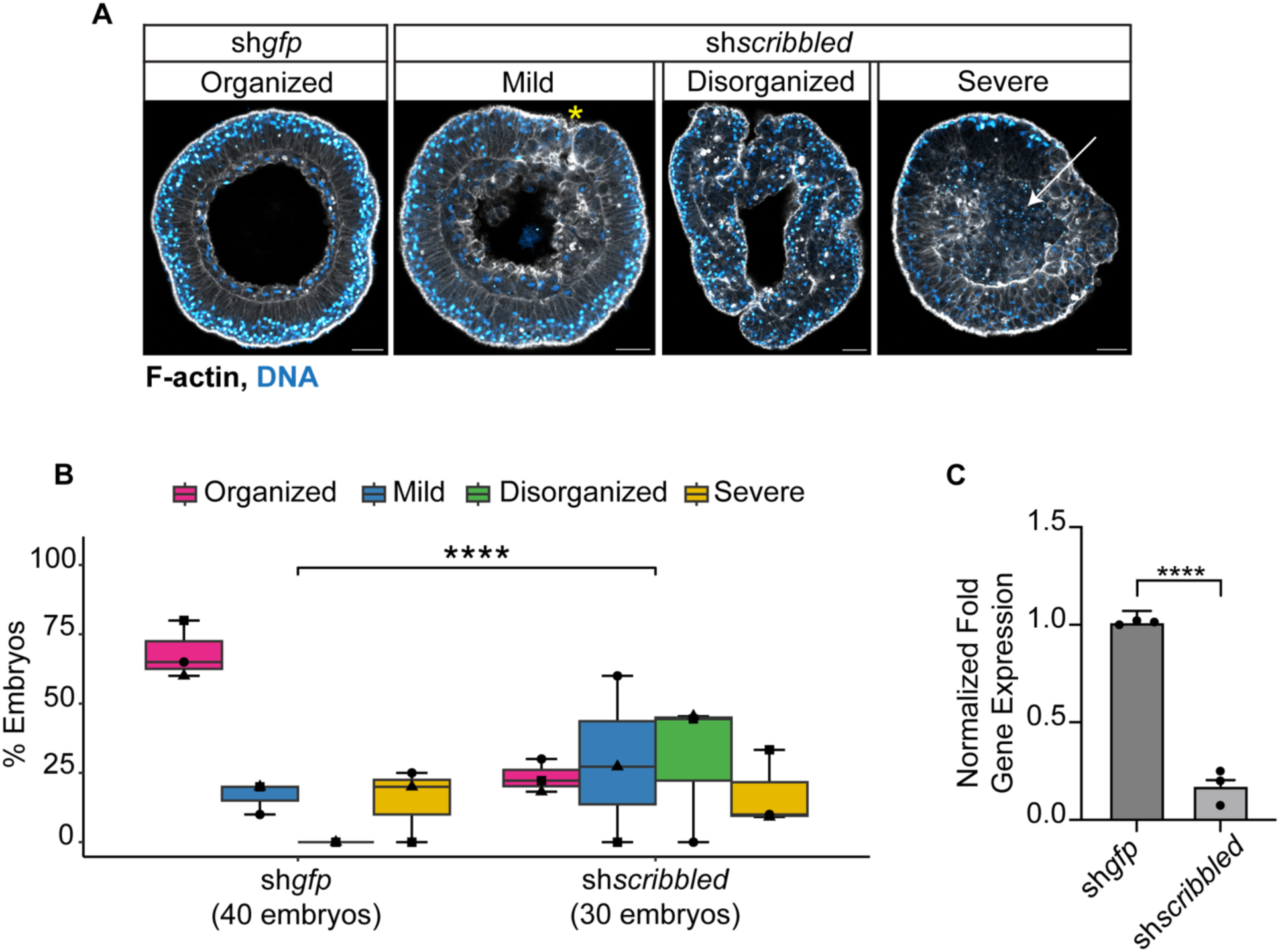
Epithelial organization is impaired in *scribbled* mutant embryos. (A) Ectodermal epithelia are disorganized in sh*scribbled* embryos and manifest surface depressions similar to sh*lgl* embryos. (B) Quantification of observed phenotypes similar to those described in Figure 2. “Mild” refers to embryos with disorganization in restricted areas of the ectoderm, shown near the yellow star in “mild” embryo of (A). ”Disorganized” refers to embryos manifesting global disorganization with depressions and misshapen cells throughout the ectoderm. “Severe” refers to embryos undergoing cell lysis pointed by the white arrow as an obliterated cell mass in “severe” embryo of (A). Fisher’s exact test; ****p< 0.0001). Symbols ▪▴● correspond to embryo percentages from three independent experiments. (C) Expression of *scribbled* is reduced in post-gastrula (28hpf) sh*scribbled* embryos compared to sh*gfp* controls, as verified by qPCR of mRNA levels from embryos of three independent experiments. (Wilcoxon rank sum test; ****p< 0.0001). Images show mid-sagittal sections from post-gastrula embryos. Scale bars: 20μm.

**Fig S3.**
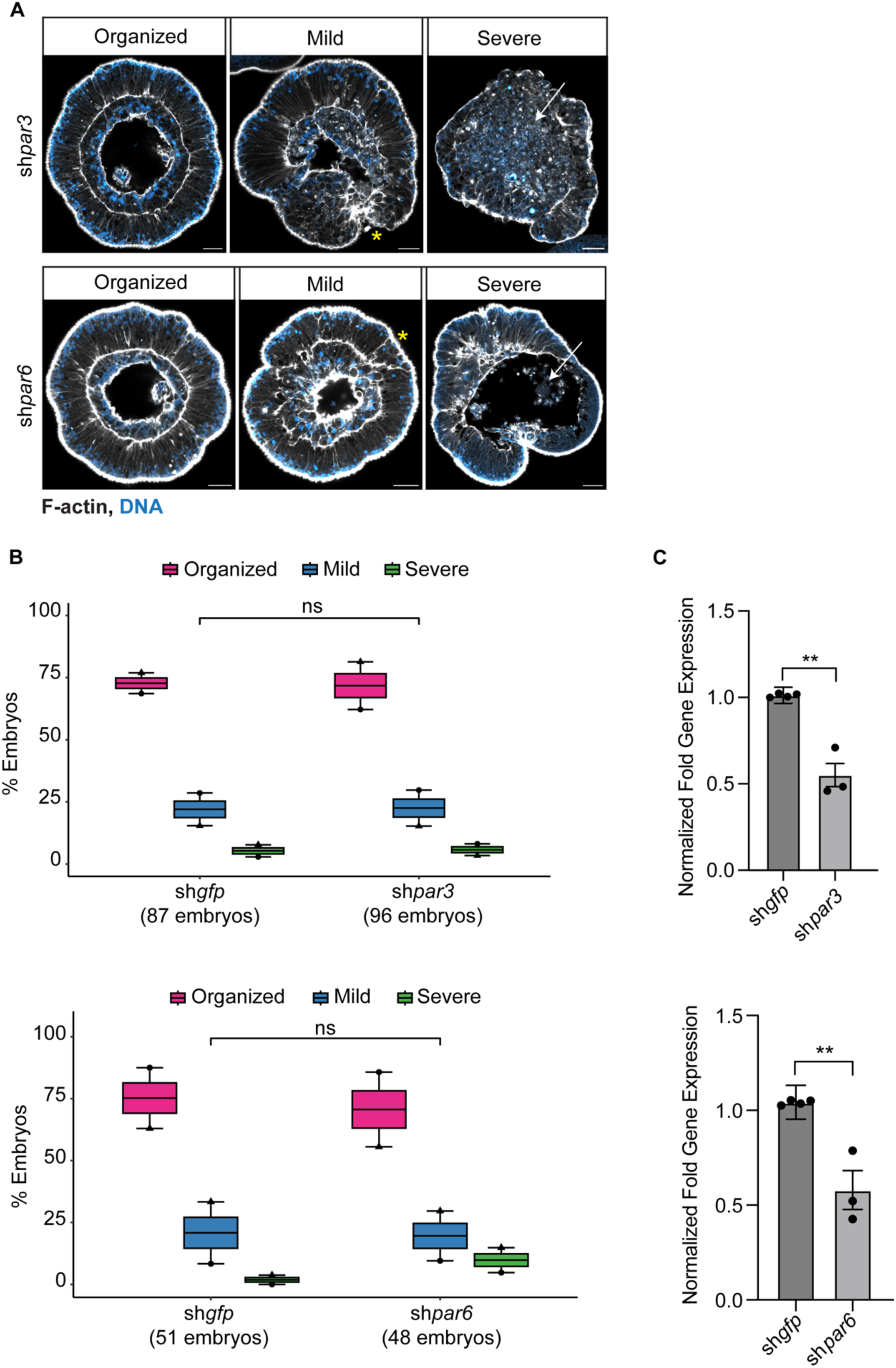
Reduced expression of apical *par3 and par6* does not compromise epithelial integrity. (A) Embryos with reduced *par3* and *par6* expression develop normal ectodermal epithelia. (B) Quantification of observed phenotypes from two independent experiments. “Mild” refers to embryos with ectodermal epithelial disorganization in restricted areas of the ectoderm, shown near the yellow asterisk in “mild” embryos of (A). “Severe” refers to embryos undergoing cell lysis pointed by the white arrows as an obliterated cell mass in “severe” embryos of (A). (Fisher’s exact test; ns p> 0.05). Symbols ▴● correspond to embryo percentages from two independent experiments. (C) Expression of *par3* and *par6* is reduced in post-gastrula (28hpf) sh*par3 and* sh*par6* embryos compared to sh*gfp* controls, as verified by qPCR of mRNA levels from embryos of three independent experiments. (Wilcoxon rank sum test; **p< 0.01). Images show mid-sagittal sections from post-gastrula embryos. Scale bars: 20μm.

**Fig S4.**
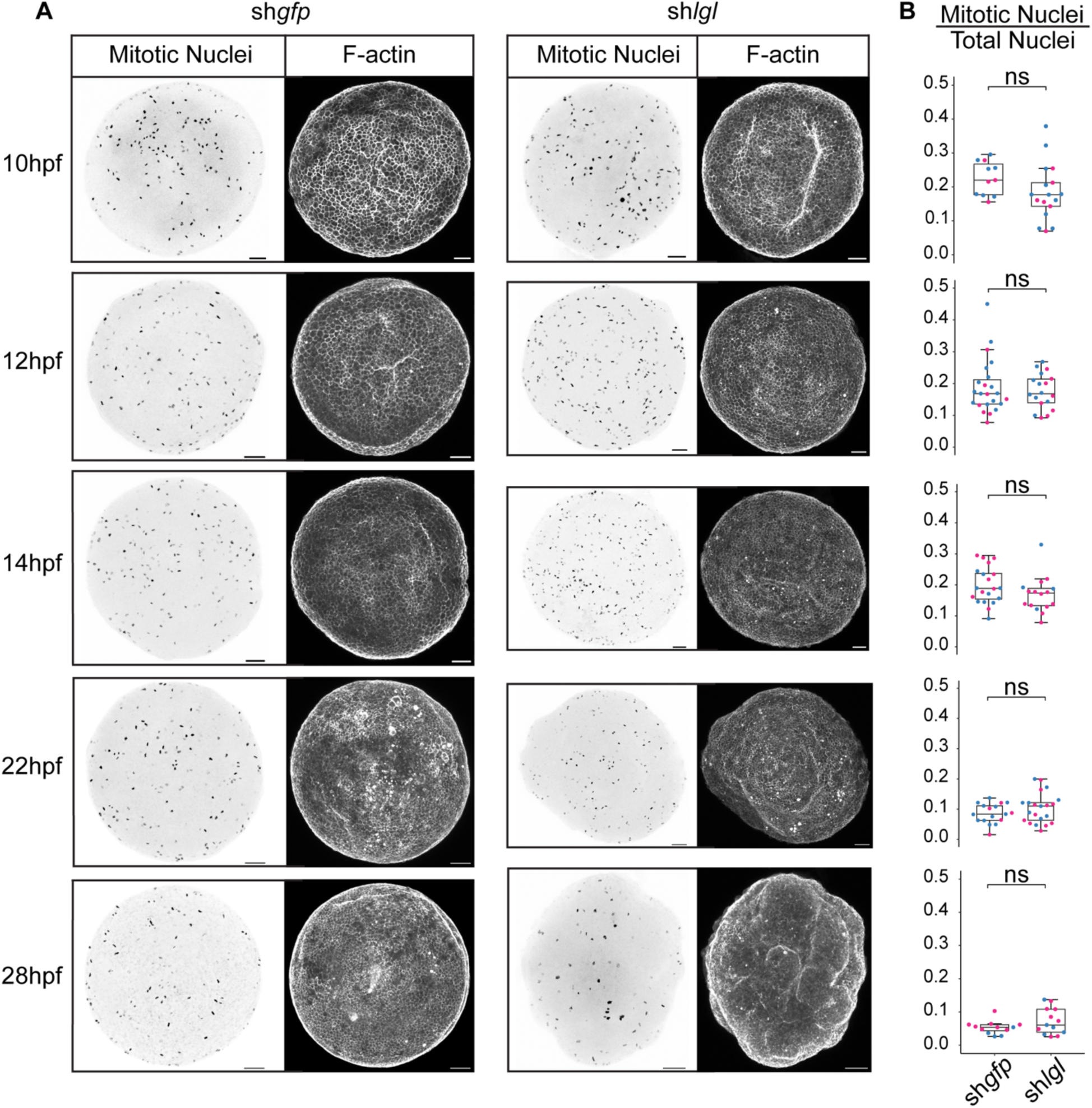
*lgl* mutant embryos do not exhibit cell over proliferation during development. (A) Mitotic nuclei stained with PH3 antibody and corresponding F-actin-stained sh*gfp* control and sh*lgl* embryos across the indicated time points during development. (B) Quantification of ratios of mitotic nuclei / total nuclei numbers in sh*gfp* control and sh*lgl* embryos across the corresponding times during development. Magenta and blue dots correspond to average values from each embryo analyzed from two independent experiments. Number of sh*gfp* embryos analyzed per time point: 10hpf (11 embryos), 12hpf (23 embryos), 14hpf (20 embryos), 22hpf (17 embryos) and 28hpf (12 embryos). Number of sh*lgl* embryos analyzed per time point: 10hpf (17 embryos), 12hpf (18 embryos), 14hpf (16 embryos), 22hpf (21 embryos) and 28hpf (13 embryos). (Wilcoxon rank sum test; ns p> 0.05). Images show projections of whole embryos. Scale bars: 20μm.

**Fig S5.**
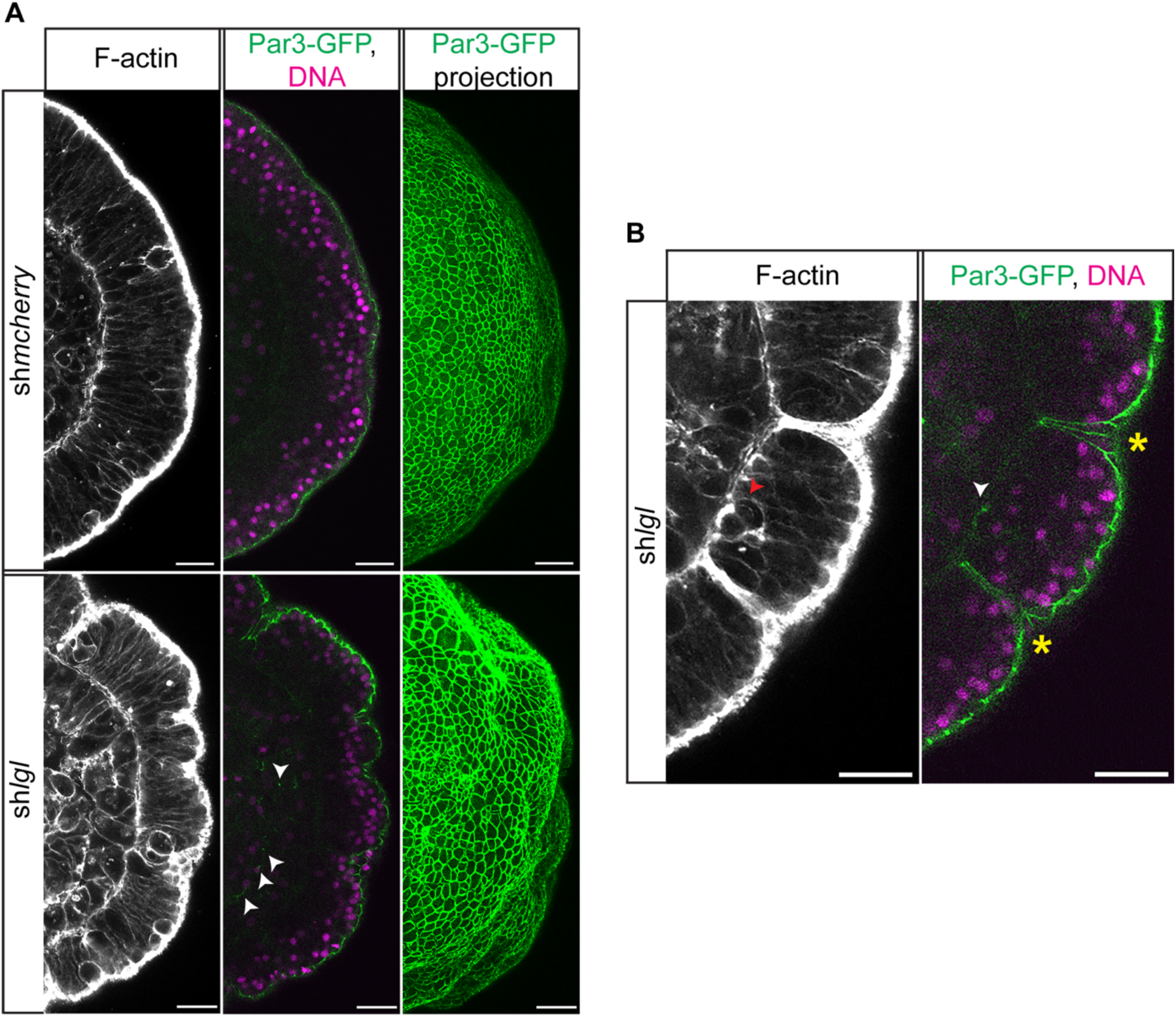
Par3-GFP is mis-localized in the basal region of the *lgl* mutant ectodermal epithelia. (A) Par3-GFP is normally enriched exclusively in the apical cell junctions, as shown in the mid-sagittal section and surface projection views of sh*mcherry* control embryos. In sh*lgl* embryos, Par3-GFP is enriched in apical cell junctions, similar to controls, as well as in some basal junctions of the ectodermal epithelium pointed by the white arrowheads. (B) Closeup into a region of the sh*lgl* ectodermal epithelium. Par3-GFP is enriched in apical and some basal junctions of the epithelium pointed by the white arrowhead. Basal cells with Par3-GFP enrichment are round, as shown in the F-actin labelled mid-sagittal section of the same region (red arrow). Par3-GFP is also found along the surface of cells lining the ectodermal depressions pointed by the yellow stars. All images show post-gastrula embryos. Scale bars: 20 μm.

**Fig S6.**
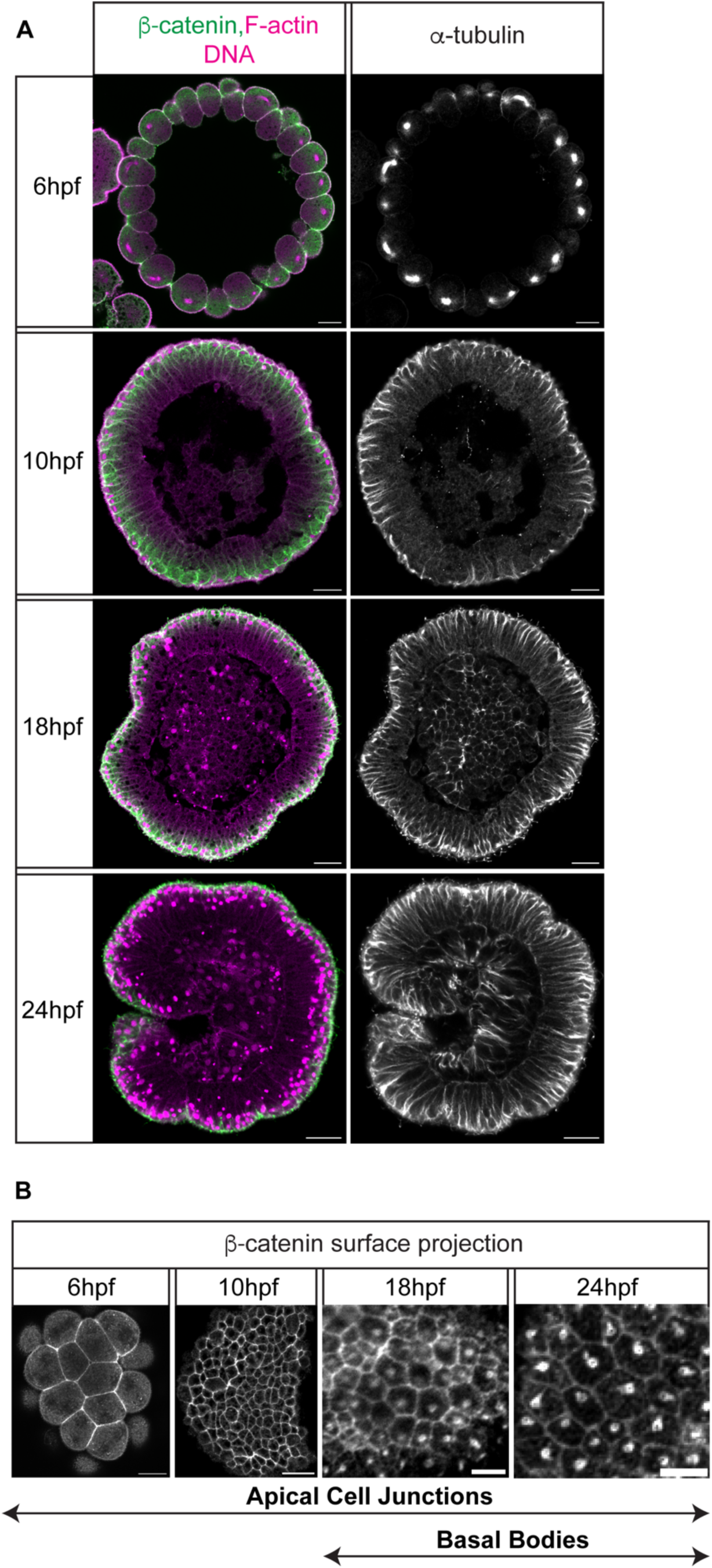
β-catenin is enriched at cell junctions and the basal body in *Nematostella* ectodermal epithelia. (A) mid-sagittal and (B) surface views of β-catenin localization in developing embryos. β-catenin is enriched at apical cell junctions throughout development (6-24hpf). β-catenin is also enriched at the basal bodies that become evident during late blastula and gastrula stages (18-24hpf). During these stages cilia form on the embryo surface as marked with α-tubulin in 18hpf and 24hpf embryos (A). Scale bars: 20μm (A and B for 6hpf and 10hpf embryos) and 5μm (B for 18hpf and 24hpf embryos).

**TABLE S1:**
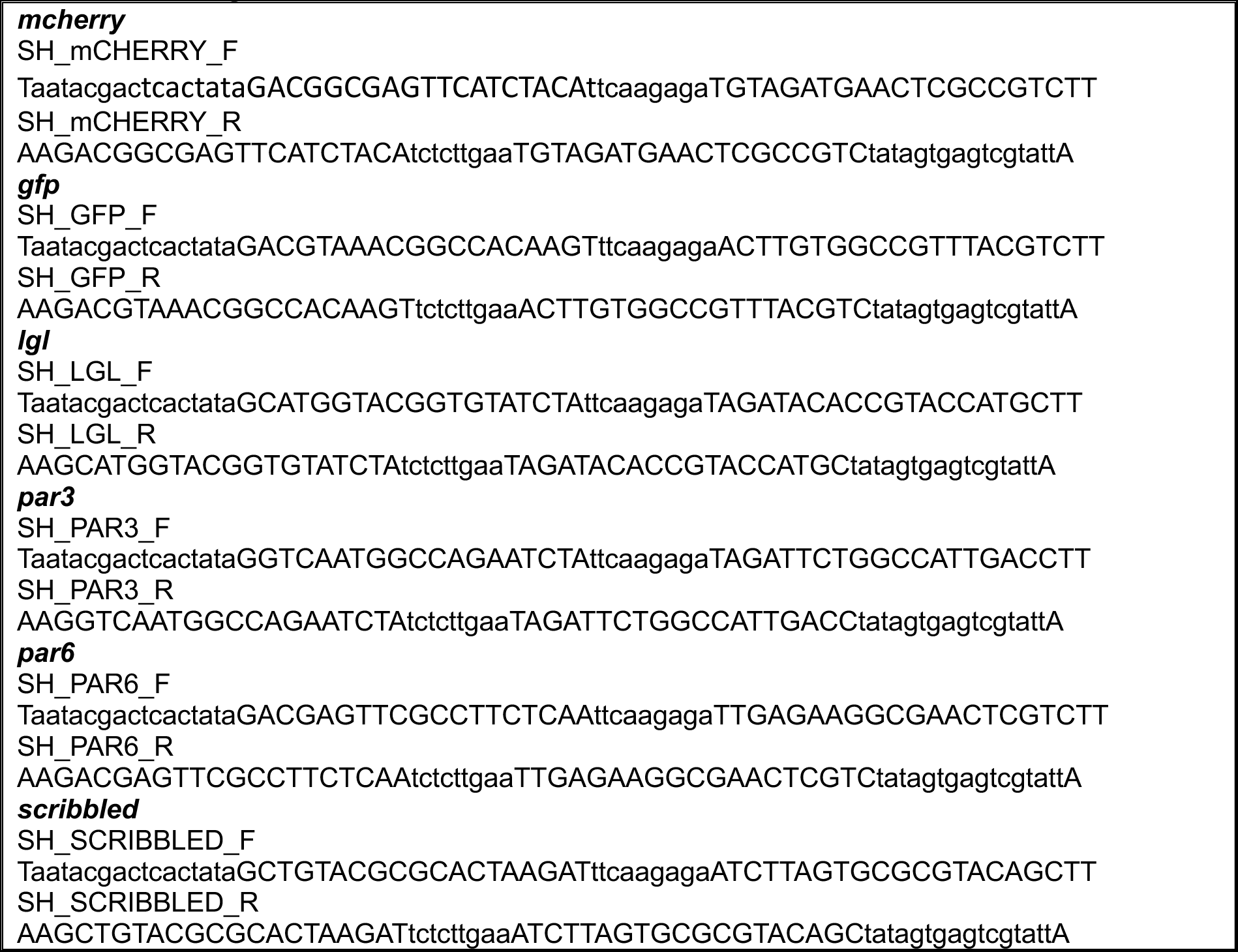
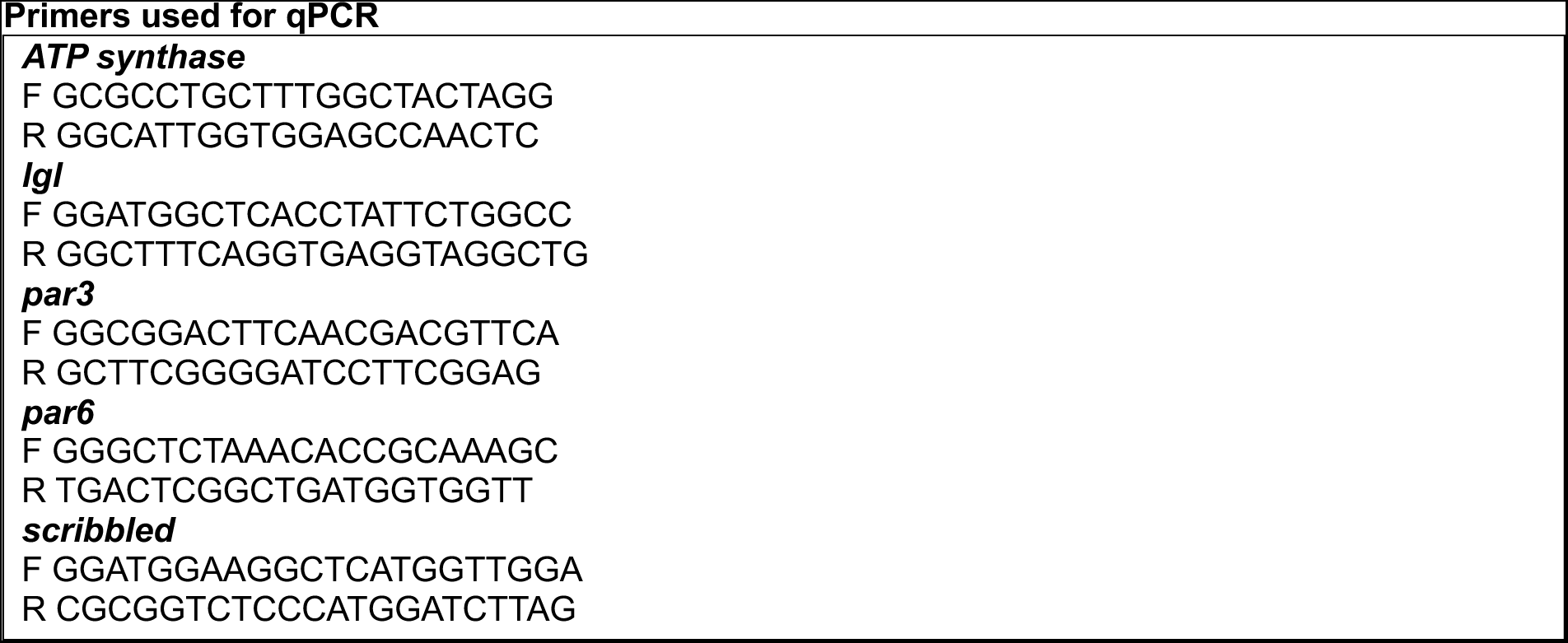
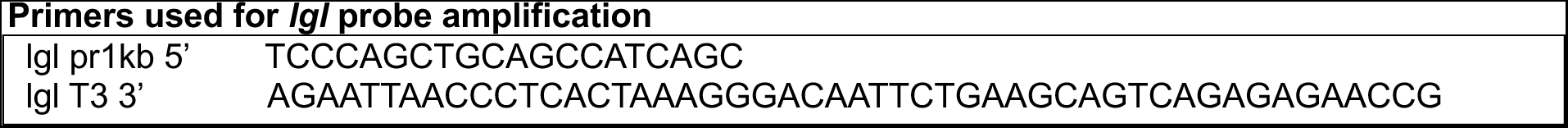
Primers used for gene knockdown.

